# Multilayered extracellular matrix–derived scaffolds direct progenitor cell differentiation *in vitro* and osteochondral-tissue formation *in vivo*

**DOI:** 10.64898/2026.08.28.747815

**Authors:** Giovanni Gonnella, Olivia Strong, Vasile M. Sularea, Aliaa S. Karam, Gabriela S. Kronemberger, Daniel J. Kelly

**Affiliations:** Trinity Centre for Biomedical Engineering, Trinity Biomedical Sciences Institute, Trinity College Dublin, Dublin, Ireland; Department of Mechanical, Manufacturing and Biomedical Engineering, School of Engineering, Trinity College Dublin, Dublin, Ireland; Department of Anatomy and Regenerative Medicine, Royal College of Surgeons in Ireland, Dublin, Ireland; Advanced Materials and Bioengineering Research Centre (AMBER), Royal College of Surgeons in Ireland and Trinity College Dublin, Dublin, Ireland; School of Biochemistry and Immunology, Trinity Biomedical Sciences Institute, Trinity College Dublin, Dublin, Ireland

**Keywords:** extracellular matrix, osteochondral repair, multilayer scaffold, articular cartilage, subchondral bone, caprine model

## Abstract

Osteochondral repair requires restoration of zonally organised articular cartilage and subchondral bone, yet translatable implants rarely reproduce this spatial complexity. Here, we developed an acellular multilayer scaffold comprising a superficial 2% (w/v) articular cartilage extracellular matrix (AC-ECM) phase, an intermediate 5% AC-ECM phase and a basal 6% bone ECM (BN-ECM) phase. The scaffold formed continuous interfaces, displayed region-dependent pore architecture and showed limited residual deformation after compression. In vitro, constructs seeded with caprine mesenchymal stromal and articular cartilage progenitor cells supported cell expansion and the accumulation of sulfated glycosaminoglycan- and collagen-rich matrix, with regional differences in collagen I, II and X immunoreactivity. Following eight weeks of subcutaneous implantation, cell-seeded scaffolds contained more collagenous matrix than unseeded controls, while vascularisation preferentially localised to the BN-ECM phase. In a six-month caprine osteochondral defect model, scaffold treatment significantly improved macroscopic and histological repair and increased PTA-attenuating, collagen-rich repair-tissue fill within the chondral region (∼60% versus ∼40%). It also limited extension of this tissue into the subchondral region and produced superficial collagen fibres more closely aligned parallel to the articular surface. Repair tissue exhibited greater collagen II immunoreactivity, increased *ACAN* and *COL2A1* expression and reduced *COL10A1* expression, whereas deeper mineralised tissue formation was not significantly improved. These findings demonstrate that this acellular multilayer ECM scaffold improves the cartilage component of osteochondral repair without exogenous cells or growth factors and identify subchondral bone regeneration as the principal remaining design challenge.

## 1. Introduction

Articular cartilage (AC) is a highly specialised connective tissue whose load-bearing function depends on a depth-dependent collagen architecture and a proteoglycan-rich extracellular matrix (ECM), which together function to distribute mechanical loads across synovial joint surfaces [1,2]. Because AC is avascular and has a limited intrinsic capacity for repair, focal injuries frequently result in the formation of a mechanically inferior fibrocartilaginous tissue and may contribute to the development of osteoarthritis [3–5]. Advanced joint degeneration can ultimately require total joint arthroplasty, in which the damaged osteochondral tissues are replaced using permanent metallic and polymeric implants [6]. There is therefore a need for regenerative strategies capable of restoring both the articular cartilage and the underlying tissues that support its long-term function.

Current cartilage and osteochondral repair procedures each present important limitations. Osteochondral autograft and allograft transplantation can provide immediate replacement of damaged tissue with hyaline cartilage supported by subchondral bone [7,8]. However, autografts are constrained by donor-site morbidity and the limited amount of tissue available, while allografts are affected by donor availability, cost and procurement- and processing-related risks [7–9]. Several multiphasic osteochondral scaffolds have also reached clinical evaluation and incorporate materials including collagen, hyaluronic acid, synthetic polymers and mineral components [10–12]. Nevertheless, these constructs often lack the tissue-specific biochemical complexity and depth-dependent organisation of the native osteochondral unit. This may limit their ability to direct endogenous stem and progenitor cells towards the distinct phenotypes required for stable cartilage, interface and bone regeneration.

Successful osteochondral repair requires more than the regeneration of cartilage or bone in isolation. The osteochondral unit is a hierarchically organised structure comprising superficial and deep articular cartilage, a calcified cartilage interface and the underlying subchondral bone. These regions possess distinct cellular, biochemical, structural and mechanical properties, while remaining mechanically and biologically interconnected [1,2,13,14]. In particular, the transition from an avascular, proteoglycan-rich cartilaginous matrix to a vascularised and mineralised bone environment occurs across a relatively narrow interface. A homogeneous scaffold is therefore unlikely to provide all the regional signals required to maintain a stable cartilaginous phenotype at the articular surface while simultaneously supporting vascularisation and spatially defined mineralised tissue formation at depth.

Tissue-derived extracellular matrix (ECM) scaffolds offer a potential means of addressing this limitation. When appropriately decellularised and processed, ECM-derived biomaterials can retain structural proteins, glycosaminoglycans, adhesive ligands and matrix-bound signalling molecules capable of influencing cell recruitment, differentiation and matrix deposition [15–19]. The biological activity of an ECM scaffold depends not only on its tissue of origin but also on its processing conditions, matrix concentration, pore architecture and mechanical properties. Cartilage- and bone-derived ECMs provide distinct compositional environments associated with their respective tissues and can therefore deliver tissue-relevant biochemical signals to resident cells [17,20–24], including immune cells that define the inflammatory and regenerative environment and host stem/progenitor cells that may be capable of generating functional repair tissues [25,26]. Spatially organising these tissue-specific matrices within a single construct may consequently generate regionally distinct microenvironments that better support osteochondral tissue formation.

Multilayer scaffold design offers one potential strategy for recreating the depth-dependent heterogeneity of the native osteochondral unit [27,28]. Previous work from our group demonstrated that bilayered ECM-derived scaffolds could enhance osteochondral defect repair by providing spatially distinct environments supportive of chondrogenic and osteogenic tissue formation [29]. More broadly, the spatial organisation of cartilage- and bone-associated ECMs within multiphasic constructs has been shown to influence cell phenotype and tissue formation through the scaffold depth [26,29]. However, variability in the quality and spatial organisation of the repair tissue generated using bilayered scaffolds indicated that further refinement of the composition and architecture of the constructs was required. In particular, a two-layer configuration does not explicitly reproduce the transitional environment separating the superficial cartilage and underlying subchondral bone compartments.

In addition to ECM source, matrix concentration provides a means of modulating scaffold architecture and mechanical behaviour. Increasing ECM concentration generally increases matrix density and stiffness while also influencing pore architecture, transport properties and the amount or accessibility of tissue-derived biochemical cues. Matrix stiffness is an important regulator of stem and progenitor cell attachment, spreading, differentiation and matrix production through mechanisms that include cytoskeletal tension and mechanotransduction [14,30–32]. However, changing ECM concentration simultaneously alters multiple structural, mechanical and compositional properties, and its biological effects cannot therefore be attributed to stiffness alone. Instead, concentration can be considered a composite scaffold-design parameter through which the biochemical and biophysical environment presented to resident cells may be adjusted [14,31,33].

Based on this rationale, we developed a tri-layer ECM-derived scaffold in which both ECM source and concentration were varied through the construct depth. The scaffold comprised a low-concentration 2% AC-ECM superficial region, a higher-concentration 5% AC-ECM middle or transition region and a 6% bone-derived ECM (BN-ECM) bottom region. The superficial AC-ECM region was intended to provide a permissive environment for cartilage-like matrix formation, based on our recent findings that 2% AC-ECM supports a hyaline cartilage phenotype [34]. The denser middle AC-ECM region was introduced to provide a transitional microenvironment representative of deeper cartilage and the osteochondral interface, based on our recent findings that 5% AC-ECM supports the development of a more hypertrophic cartilage phenotype [34]. The bottom BN-ECM region was incorporated to provide increased matrix density, mechanical support and bone-associated compositional cues, again based on our recent findings such stiffer scaffolds preferentially support an osteogenic phenotype [35].

The present design therefore differs from the previous bilayered scaffold by incorporating three, rather than two, distinct regions and by varying ECM concentration in addition to tissue source. This approach was intended to produce a more gradual transition between the cartilage- and bone-associated environments and provide resident cells with depth-dependent mechanical [14,30–33] and compositional signals [17,20–24]. The three regions were integrated during fabrication using a controlled lyophilisation protocol designed to generate a continuous porous construct, increase pore size, while minimising the risk of separation between independently manufactured layers [29,36].

We hypothesised that spatially modulating both ECM source and concentration through the scaffold depth would generate distinct but continuous microenvironments capable of supporting regionally appropriate cell phenotypes and osteochondral tissue formation. To test this hypothesis, scaffold architecture, pore structure and compressive behaviour were first characterised relative to the corresponding single-phase formulations. The capacity of the multilayer construct to support cell proliferation and spatial matrix deposition was subsequently evaluated *in vitro* by seeding these scaffolds with mesenchymal stem/stromal cells (MSCs) and articular cartilage progenitor cells (ACPs). Cell-seeded scaffolds were then implanted subcutaneously in nude mice to determine whether regional patterns of matrix formation and vascularisation were maintained *in vivo*. Finally, to assess their translational potential, acellular multi-layered ECM scaffolds were implanted for six months in a clinically relevant caprine osteochondral defect model to evaluate their capacity to recruit endogenous cells and promote tissue repair under physiological joint loading.

## 2. Materials and methods

### 2.1. Articular cartilage ECM extraction

Articular cartilage (AC) was harvested from the condyles of 6-month-old porcine donors obtained from a local abattoir following a previously established protocol [29,37]. Briefly, articular cartilage was removed from the tibia and femur using 6 mm biopsy punches and finely diced into small pieces (3–4 mm) using a commercial knife. The tissue was weighed, transferred into 50 mL conical tubes, and pre-treated with 0.2 M NaOH for 24 hours at 4 °C to remove non-specific proteins and sulfated glycosaminoglycans (sGAGs) [38,39]. After pre-treatment, the NaOH solution was removed, and the cartilage was thoroughly washed with ultrapure water. The extracellular matrix (ECM) was then enzymatically extracted using a solution of 1500 U/mL of pepsin (Sigma) in 0.5 M acetic acid (HAc, Sigma), under slow rotation at 4 °C for 24 hours. The tubes were centrifuged at 2500 g for 1 hour at 4 °C to remove non-solubilized material, with the pellet discarded and the supernatant retained. The supernatant, containing solubilized ECM, was transferred to new conical tubes, and collagen type II fibers were salted out by adding 5 M sodium chloride (NaCl, Sigma) to a final concentration of 0.9 M NaCl overnight at 4 °C [40]. The tubes were centrifuged at 2500 g for 1 hour at 4 °C, and the supernatant was discarded. The collagen pellet was resuspended overnight in 0.5 M HAc at 4 °C. The salt precipitation step was repeated once more to increase ECM purity. The solubilized ECM was then transferred to 6–8 kDa dialysis membranes (Spectrum Labs) and dialyzed against ultrapure water for 48 hours, with the solution changed every 24 hours. Finally, the ECM was transferred into petri dishes and lyophilized using a Labconco (FreeZone Triad, Labconco, KC, USA) freeze-drier. The lyophilized ECM was stored at -20 °C until further use.

### 2.2. Bone ECM extraction

Bone-ECM (BN-ECM) was extracted from 6-month-old porcine femoral heads obtained from a local abattoir by adapting previously described methods [22,41–43]. Briefly, cryomilled bone powder was transferred into 50□mL Falcon tubes (Sigma) and subjected to an initial pre-treatment consisting of incubation in 0.1□M sodium hydroxide (NaOH, Sigma) at 4 °C for 48 hours, followed by 10 % (v/v) 1-butanol (Sigma) at 4 °C for an additional 48 hours. Both solutions were refreshed every 12 hours. This sequential treatment is performed to effectively remove non-specific proteins, sulphated glycosaminoglycans (sGAG) and lipids, which can interfere with ECM extraction and reduce final product quality. Following pre-treatment, bone powder was thoroughly washed in phosphate-buffered saline (PBS), and the ECM was subsequently enzymatically extracted using 1500□ U/mL pepsin (Sigma) in 0.5□M acetic acid (HAc, Sigma). This step was conducted under slow rotation at 4 °C for 72 hours. Insoluble material was removed by centrifugation at 3000□×□g for 90 minutes at 4 °C, and the resulting supernatant was transferred to fresh Falcon tubes while the pellet was discarded. To precipitate collagen fibres, 5□M sodium chloride (NaCl, Sigma) was added to the supernatant to reach a final concentration of 2□M. The solution was mixed gently and left to precipitate overnight at 4 °C. After precipitation, samples were centrifuged at 3000□×□g for 90 minutes at 4 °C. The supernatant was discarded, and the collagen-rich ECM was resuspended in 0.5□M acetic acid by slow rotation at 4□rpm overnight at 4 °C to avoid collagen degradation. The salt-precipitation step was then repeated once more to enhance ECM purity. The acid-solubilised ECM solution was transferred into dialysis membranes (molecular weight cut-off: 6–8□kDa; Spectrum Labs) and dialysed against 0.1□M acetic acid for 48 hours at 4 °C, with a complete solution change after 24 hours. Following dialysis, the ECM solution was poured into sterile Petri dishes (Sigma) and lyophilised using a Labconco freeze-dryer (FreeZone Triad, Labconco, KC, USA). The lyophilised ECM was stored at −20 °C until further use in scaffold fabrication.

### 2.3. Multilayer scaffold fabrication

Tissue-specific multilayer scaffolds were fabricated from three solubilised ECM formulations: 2% AC-ECM, 5% AC-ECM and 6% BN-ECM. The method was adapted from *Browe et al.* [27]. Following partial crosslinking with 5 mM glyoxal at 37 °C for 30 min, the 6% BN-ECM slurry was pipetted into a custom mould designed to promote preferential heat transfer through the metal base (polydimethylsiloxane [PDMS] wells) and frozen in liquid nitrogen. The mould was then removed from liquid nitrogen, and the 5% AC-ECM slurry was pipetted on top of the frozen BN-ECM layer and allow freezing. The 2% AC-ECM slurry was then added as the top final layer and allow freezing. The filled mould was equilibrated at room temperature for 10 min before freeze-drying using the previously described annealing protocol [27]. After lyophilisation, scaffolds were physically crosslinked by dehydrothermal treatment at 115 °C under 2 mbar vacuum for 24 h in a vacuum oven (VD23, Binder, Germany), as previously described [34].

Scaffolds for the subcutaneous study were 5 mm in diameter and 4.5 mm high, with three 1.5-mm regions. Enlarged scaffolds for the caprine study were 7 mm in diameter and 6 mm high, comprising a 3-mm BN-ECM region, a 1.5-mm middle AC-ECM region and a 1.5-mm superficial AC-ECM region.

### 2.4. Mechanical testing

The compressive properties of the multilayer scaffolds and corresponding single-phase 2% AC-ECM, 5% AC-ECM and 6% BN-ECM controls (diameter 5 mm, height 4.5 mm; n = 4 per formulation) were assessed using a Zwick materials-testing machine (Herefordshire, UK) equipped with a 5 N load cell. Constructs were kept in a PBS bath to maintain hydration and were tested at room temperature. An initial preload of 0.005 N was applied for 60 s to ensure contact with the compression plates. Force was then zeroed and samples were subjected to unconfined stress-relaxation compression at 10%, 20% and 30% strain at 2.3% strain/s, with a 30-min relaxation period after each step. After the 30% strain step, 10 compression cycles were applied at 1 Hz. Load and displacement were recorded continuously. Stress was calculated by dividing load by the initial cross-sectional area, and strain by dividing displacement by the initial sample height.

To quantify residual deformation, compression tests were performed to 10%, 20% and 30% strain, and residual deformation was recorded after unloading at each strain level.

**[AUTHOR QUERY: Confirm whether residual deformation was measured using the same or separate specimens, the recovery interval after unloading and the calculation used.]**

### 2.5. Isolation and expansion of articular cartilage progenitor cells (ACPs) and mesenchymal stromal/stem cells (MSCs)

ACPs were isolated from full-thickness AC shavings of skeletally mature female goats through differential adhesion to fibronectin. AC shavings were obtained from animals euthanized as part of a separate, ethics-approved project by the Irish Health Products Regulatory Authority (approval number AE18982). Cell culture flasks were coated with 10 μg/mL fibronectin (Brennan & Co) in 0.1 M Dulbecco’s phosphate buffer (PBS) with 1 mM magnesium chloride and 1 mM calcium chloride (CaCl_2_) overnight at 4 °C (all Sigma, Ireland) [44]. The AC was rinsed with PBS and then digested in pronase (70 U/mL, Thermo Fischer Scientific) for 30 min at 37 °C. Using a scalpel, AC shavings were chopped to 1–2 mm^3^ pieces. The minced AC was then digested in collagenase type II (300 U/mL, Gibco, Ireland) overnight at 37 °C. Enzymes were prepared in Dulbecco’s Modified Eagle Medium/Nutrient Mixture F-12 (DMEM/F-12, ThermoFischer Scientific) supplemented with 100 U/mL penicillin, 100 μg/mL streptomycin, and 2.5 μg/mL amphotericin B (all Gibco, Ireland). Next the digested AC was filtered through a 70 μm sieve and centrifuged at 500 x g for 5 min. Cells were then seeded into the fibronectin coated flasks. After 20 min, non-adherent cells (chondrocytes) were discarded while adherent cells (ACPs) were kept in culture. ACPs were cultured in XPAN-12 composed of DMEM/F-12 supplemented with 10 % (v/v) FBS, 100 U/mL penicillin, 5 ng/mL of FGF-2 (PeproTech, USA), 100 μg/mL streptomycin, and 2.5 μg/mL amphotericin B. On day 2, ACPs were switched to A_XPAN composed of XPAN-12 supplemented with 0.5 μg/mL of L-ascorbic acid 2-phosphate (Sigma, Ireland) and 1 ng/mL of TGF-β1 (PeproTech, USA). Media changes were performed every 2–3 days. At 80 % confluency, ACPs were harvested using TrypLE (Biosciences, Ireland).

Bone marrow-derived mesenchymal stromal/stem cells (MSCs) were obtained following a previously established method [22,45,46]. Briefly, MSCs were isolated from the femurs of a 4-month-old caprine donor acquired from a local veterinary (Lyon’s Farm, University College Dublin) following all relevant guidelines and regulations. The bone marrow was extracted from the femoral shaft and washed with growth medium, which consisted of high-glucose DMEM (Biosciences, Ireland) supplemented with 10% fetal bovine serum (FBS, GIBCO, Biosciences, Ireland) and 2% penicillin (100 U/mL) and streptomycin (100 µg/mL) (Biosciences, Ireland). A homogeneous suspension was prepared by triturating the marrow with a needle. The solution was centrifuged twice at 650 g for 5 minutes, with the supernatant discarded each time. The cell pellet was triturated and filtered through a 40 µm cell sieve. Following colony formation, cells were trypsinized, counted, and re-plated at a density of 5 × 10³ cells/cm² for further passage. All cell expansion was conducted under normoxic conditions in growth medium, with media changes occurring twice weekly. Cells were used at the end of passage 3.

### 2.6. Cell culture

For *in vitro* studies, caprine bone marrow mesenchymal stromal cells (MSCs) or articular cartilage progenitor cells (ACPs) were harvested and seeded at a density of 1 x 10^6^ cells per scaffold as previously described [29,47]. Briefly, in the MSC/MSC groups, MSCs were seeded on both sides of the scaffolds, while for the ACP/MSC groups, ACPs were seeded on the AC side and MSCs on the BN side, with 0.5 × 10^6^ cells per side. Before supplementing the media, the cells were allowed to attach to the scaffolds for a total of 2 h in the incubator at 37 °C (60 min-per-side).

To assess cell differentiation and tissue deposition, the constructs were cultured in chondrogenic media consisting of hgDMEM GlutaMAX supplemented with penicillin/streptomycin, sodium pyruvate (100 µg·mL−1), L-proline (40 µg·mL−1), L-ascorbic acid-2-phosphate (50 µg·mL−1), linoleic acid (4.7 µg·mL−1), bovine serum albumin (1.5 mg·mL−1), 1× insulin−transferrin−selenium (ITS), dexamethasone (100 nM) (all Sigma), and human transforming growth factor-β3 (TGF-β3, 10 ng·mL−1; Peprotech, UK).

Samples at day 0 (after cell seeding) and day 28 were biochemically analysed for DNA, sulphated GAG (sGAG), and collagen content. Quantification of dsDNA in the digested constructs was performed using a Quant-iT Pico Green dsDNA kit (Invitrogen) according to the manufacturer’s protocol. sGAG quantification was performed using a 1,9-dimethylmethylene blue (DMMB) assay according to the manufacturer’s protocol with chondroitin 4-sulfate used as a reference standard (Blyscan sulfated sGAG assay kit, Biocolor, Northern Ireland). Collagen content of scaffolds was quantified by measuring total hydroxyproline content as previously described [48,49]. A hydroxyproline:collagen ratio of 1:7.69 was assumed to determine the collagen content [50]. Furthermore, due to the scaffolds being composed predominantly of collagen, the *net collagen accumulated* data was calculated as day-28 collagen minus day-0 collagen.

For the murine subcutaneous implantation study, caprine MSCs and ACPs were seeded on the scaffolds the evening prior to surgery following the same logic as for the *in vitro* study. Constructs were maintained in expansion media: DMEM + GlutaMAX^TM^, 10 % fetal bovine serum (Gibco^®^) with 100 units/ml Penicillin, 100 units/ml Streptomycin (Gibco^®^) overnight before surgery.

### 2.7. Histological and immunohistochemical analysis

All samples were washed in PBS following overnight fixation in 4 % paraformaldehyde (PFA, Sigma). Samples were subsequently dehydrated and wax embedded. Embedded samples were then sectioned at a thickness of 7 μm using a microtome (Leica Biosystems, USA). Sections were stained with haematoxylin and eosin to examine matrix deposition, together with alcian blue and picrosirius red to examine sulphated glycosaminoglycan and collagen deposition, respectively. Lastly, samples were stained with alizarin red to detect any calcium deposition.

To identify specific collagen types, immunohistochemistry was performed for collagen type I, type II, and type X as previously described [22,29,37]. Briefly, sections were initially dewaxed and rehydrated, followed by performing antigen retrieval by incubation with chondroitinase ABC for collagen type I and type II, or with pronase for collagen type X. After blocking non-specific binding, sections were incubated with primary antibody overnight at 4 °C (anti-collagen type I, 1:400, type II, 1:100, and type X, 1:200, Abcam, UK). Endogenous peroxidase activity was blocked with hydrogen peroxide prior to incubation with anti-mouse IgG secondary antibody. Lastly, sections were incubated with 3,3’-diaminobenzidine (DAB) peroxidase substrate (Vector Labs, UK) to visualise positive staining.

### 2.8. Murine subcutaneous implantation

All animal experiments were conducted in accordance with the recommendations and guidelines of The Health Products Regulatory Authority (HPRA), the competent authority in Ireland responsible for the implementation of Directive 2010/63/EU on the protection of animals used for scientific purposes in accordance with the requirements of the Statutory Instrument No. 543 of 2012.

Mouse procedures were ethically approved by the Animal Research Ethics Committee of Trinity College Dublin and the HPRA under protocol AE19136/P069. Eight-week-old BALB/c OlaHsd-Foxn1nu female nude mice (Envigo, Oxford, UK) were anaesthetised with isoflurane (4% induction; 1–2% maintenance). A total of eight mice was used. Two incisions were made slightly lateral to the spine and subcutaneous pockets were created. For the first two mice, two pockets were created; up to four pockets were created in each subsequent mouse. MSC/MSC, ACP/MSC and unseeded control constructs were distributed among the pockets using randomised allocation, yielding n = 8 constructs per group across the cohort. To maintain construct shape, each scaffold was placed within a cylindrical 3D-printed resin sheath (Bio-Med Clear, LiqCreate; wall thickness 0.25 mm, diameter 5.5 mm, height 5 mm). After eight weeks, the animals were euthanised by CO□ asphyxiation and the explants were harvested. Gross morphological images were acquired before all explants underwent microcomputed tomography (µCT; n = 8 per group) at 55 kVp, 145 µA and a 10 µm voxel size using a MicroCT40 system (Scanco Medical). The 4.5-mm construct height was divided into three consecutive 1.5-mm axial regions of interest corresponding to the fabricated 2% AC-ECM, 5% AC-ECM and 6% BN-ECM regions. Following µCT, four explants per group were processed without fixation for biochemical analysis; the remaining four explants per group were fixed in 10% neutral-buffered formalin and processed for histology and immunohistochemistry. Histological sections were cut at 7 µm and stained with H&E and Masson’s trichrome.

### 2.9. Osteochondral caprine model

Caprine procedures were approved by both the University College Dublin Animal Research Ethics Committee (AREC-18-17) and the Irish Health Products Regulatory Authority (AE18982/P142). In total, n = 10 female mature healthy goats were used for the study.

The surgical procedure in the caprine model was carried out as previously described [27,44]. Briefly, the goats were pre-medicated with diazepam (0.4 mg/kg IV) and butorphanol (0.4 mg/kg IV). Once sedation was achieved, a lumbosacral epidural block was administered using lidocaine (2 mg/kg) and morphine (0.2 mg/kg). Following placement of an intravenous catheter, anaesthesia was induced with propofol to effect (maximum dose 4 mg/kg IV) and maintained using isoflurane in 100% oxygen, with ventilation to maintain end-tidal CO□ between 4.6 and 6 kPa. Lactated Ringer’s solution was infused at 10 mL/kg/h. The goats were placed in dorsal recumbency on a heat mat and each stifle joint was accessed using a lateral parapatellar arthrotomy. A critical-sized circular osteochondral defect, 6 mm in diameter × 6 mm deep, was created in the left and right lateral trochlear ridge of the distal femur using a hand drill, pointed and flat-ended drill bits and a depth guide. Saline irrigation (0.9% NaCl) was used during drilling to limit heat generation and remove debris. Stifle joints were randomly assigned to empty control or multilayer ECM scaffold treatment. A 7-mm-diameter, 6-mm-high acellular multilayer scaffold was compressed and press-fitted into each 6-mm-diameter × 6-mm-deep osteochondral defect before routine closure. Morphine (0.2 mg/kg IM) and meloxicam (0.5 mg/kg SC) were administered at the end of anaesthesia. The goats were housed in indoor pens during incision healing and allowed to bear full weight immediately. Meloxicam and amoxicillin/clavulanic acid (8.75 mg/kg) were administered daily for five days. Two weeks after surgery and suture removal, the animals were released to pasture for the remainder of the study. At six months, animals were euthanised by intravenous sodium pentobarbital overdose and the treated joints were harvested. Paired defects from the two stifles of each goat formed the paired comparison.

### 2.10. Macro scoring and μCT analysis

1.5 cm cubes containing the defect site were harvested from the goats using an oscillating bone saw. Before fixation, gross morphological images were taken with a digital microscope system (Ash Inspex HD 1080p) for macroscopic evaluation. Macroscopic images were blinded, randomised and subsequently scored by five expert reviewers using a previously described macroscopic scoring system as shown in supplementary Table 1 [29].

Before contrast-enhanced µCT, samples were stained with 1% phosphotungstic acid (PTA; Sigma) in 70% ethanol following previous protocols [48,49]. Samples were incubated at 4 °C for 19 h and scanned at 55 kVp, 145 µA and a 10 µm voxel size using the system described in Section 2.8. After scanning, samples were washed in 70% ethanol for 2 days.

PTA-attenuating collagen-rich repair tissue and mineralised tissue were quantified within a 6-mm-diameter cylindrical volume aligned with the original defect. The volume was divided into three predefined depth-based regions of interest: an upper chondral region comprising the top 1 mm, an interface or middle subchondral region comprising the next 2 mm and a deep osseous region comprising the bottom 3 mm. A greyscale threshold of 210 was applied during segmentation. For mineralised tissue, this threshold corresponds to a hydroxyapatite-equivalent density of 399.5 mg HA/cm³; this calibration value was not used to assign cartilage identity to PTA-stained soft tissue.

### 2.11. Histological and immunohistochemical analysis

For histological analysis, samples were fixed in 10% neutral-buffered formalin (Sigma) for 96 h with agitation at 4 °C. Samples were decalcified using Decalcifying Solution-Lite (Sigma) according to the manufacturer’s protocol until mineral removal was confirmed by chemical and µCT analysis. Demineralised, wax-embedded constructs were sectioned at 10 µm and stained with H&E, Safranin O, and Picrosirius Red. Histological repair was scored using a modified ICRS II system [50]. Safranin O staining was also analysed using ImageJ to quantify positively stained repair tissue within the defect. Picrosirius Red-stained sections were imaged under polarised light microscopy to assess collagen fibre orientation. Images were normalised to the local articular-surface angle, with 0° defined as parallel to the surface; axial orientations were folded to the 0–90° range. The OrientationJ plugin for ImageJ [51] was used to quantify mean fibre orientation and angular dispersion in the superficial and deep zones of the regenerated articular cartilage, as previously described [27,44]. Angular dispersion quantified the spread of fibre orientations around the mean orientation. Lower values indicate more coherent fibre alignment, whereas higher values indicate greater orientational heterogeneity. Immunohistochemistry for collagen types I, II and X was performed as described in *Section 2.7* [22,27,34].

### 2.12. Real time-qPCR analysis

Articular cartilage pieces were isolated from scaffold-treated and paired empty defects (n = 4 per group). Total RNA was isolated using the High Pure RNA Isolation Kit (Roche), and RNA concentration and purity were assessed using a NanoDrop 2000c UV–Vis spectrophotometer. RNA input was equalised before reverse transcription using the Applied Biosystems High-Capacity cDNA Reverse Transcription Kit. Quantitative real-time PCR was performed on a Bio-Rad real-time PCR detection system using TaqMan Fast Universal PCR Master Mix (Applied Biosystems) and predesigned TaqMan gene-expression assays. Expression of *COL2A1*, *ACAN*, *MMP13* and *COL10A1* was normalised to 18S ribosomal RNA. Relative expression was calculated using the 2^−ΔΔCt method with the empty group as calibrator.

### 2.13. Statistical analysis

Results are presented as mean +/- standard deviation unless otherwise specified. Experiments containing multiple formulations, regions or time points were analysed using two-way ANOVA with Tukey’s post hoc test. Paired empty and scaffold-treated defects from the caprine study were compared using paired two-tailed t-test. Statistical significance is denoted by * p < 0.05, ** p < 0.01, *** p < 0.001 and **** p < 0.0001.

## 3. Results

### 3.1. Fabrication of highly elastic multi-layered ECM scaffolds

Multilayer scaffolds were fabricated by sequentially freezing 6% BN-ECM, 5% AC-ECM and 2% AC-ECM slurries within a single mould, followed by annealed freeze-drying and DHT crosslinking (Figure 1a). SEM imaging revealed a continuous highly porous architecture throughout the depth of the multilayered scaffold, with no visible gaps or sharply defined boundaries between regions (Figure 1b). Although pore orientation was not quantitatively assessed, elongated pores showed an apparent preferential through-thickness orientation in the BN-ECM region. This directional morphology became progressively less evident toward the AC-ECM region, where pores appeared more heterogeneously oriented. Mean pore diameter was approximately 100 µm throughout the construct with slightly larger pores in the 6% BN-ECM region (∼110 µm) compared with the middle 5% AC-ECM region (∼90 µm) (Fig. 1c).

**Figure 1.**
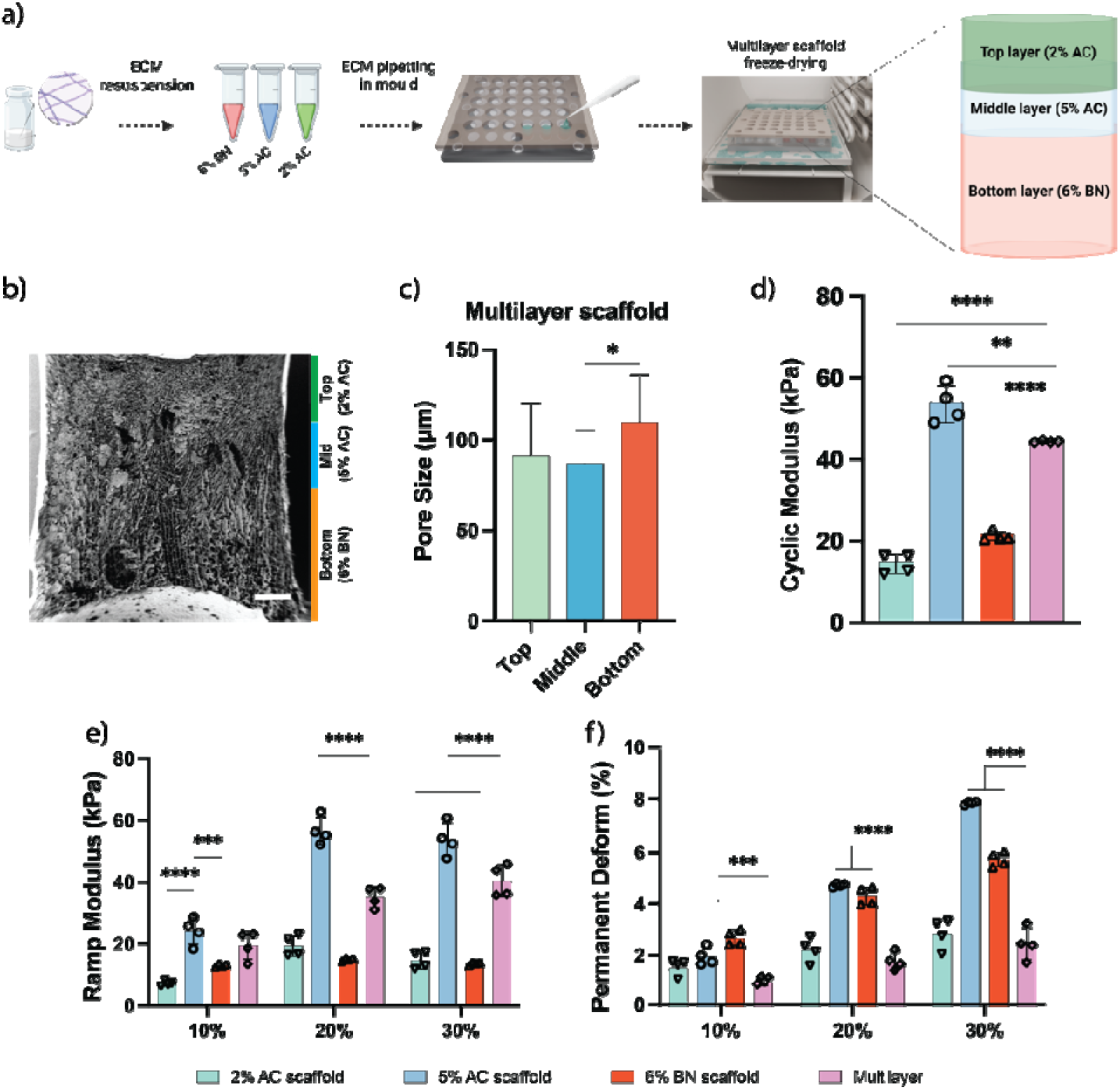
Fabrication, microarchitecture and mechanical characterisation of multilayer ECM scaffolds. (a) Schematic of sequential fabrication of multilayer scaffolds comprising a 2% AC-ECM top region, a 5% AC-ECM middle region and a 6% BN-ECM bottom region, followed by lyophilisation. (b) Representative SEM image showing continuous porous architecture across the three regions. (c) Mean pore diameter in each region. (d) Cyclic compressive modulus. (e) Ramp modulus at 10%, 20% and 30% strain. (f) Residual deformation after compression to 10%, 20% and 30% strain. Scale bar = 1 mm. Data are mean ± SD; n = 4 independent scaffolds per formulation. For pore-size analysis, five measurements were averaged for each scaffold, giving a biological n = 4. * p < 0.05, ** p < 0.01, *** p < 0.001, **** p < 0.0001.

The multilayer scaffold exhibited intermediate mechanical properties relative to the corresponding single-phase formulations. Both the ramp and cyclic moduli of the multilayer scaffold were significantly higher than the 2% AC-ECM and 5% AC-ECM scaffolds, but significantly lower than the 6% BN-ECM scaffolds (Fig. 1d, e). The multilayered scaffolds were also found to be highly elastic; following the application of 30% compression, the residual permanent deformation was approximately 2%, compared with almost 8% and 6% for the 5% AC-ECM and 6% BN-ECM scaffolds, respectively. No significant difference in permeant deformation was detected relative to the 2% AC-ECM scaffold (Fig. 1f).

### 3.2. Multilayer tissue-specific ECM scaffolds support region-dependent matrix deposition in vitro

We next sought to assess the capacity of the multi-layered scaffolds to support spatially defined chondrogenesis and osteogenesis *in vitro*. To this end the multilayered scaffolds either seeded on both sides with MSCs (termed the ‘MSC/MSC’ group), on were seeded with ACPs on their top surface and MSCs on their bottom surface (termed the ‘ACP/MSC’ group), see Figure 2a. Following 28 days of *in vitro* culture, the multilayer tissue-specific ECM scaffolds supported matrix deposition throughout the construct (Figure 2b). H&E staining confirmed the presence of cells throughout the scaffold depth (Figure 2b). Histological staining revealed some region-dependent differences in matrix deposition, with more intense staining with Alcian Blue for glycosaminoglycans evident in the top cartilage region in both the MSC/MSC and ACP/MSC groups (top layer, Figure 2b). Relatively weak staining with Alizarin Red for calcium deposition was observed throughout the depth of the scaffolds (Figure 2b). Biochemical analysis further showed increases in total DNA, sGAG, and collagen content at day 28 relative to day 0. Compared with MSC-only scaffolds, ACP/MSC co-culture constructs exhibited higher DNA at day 28 (Figure 2c), while no significant difference in sGAG and collagen deposition was observed (figure 2d-f).

**Figure 2.**
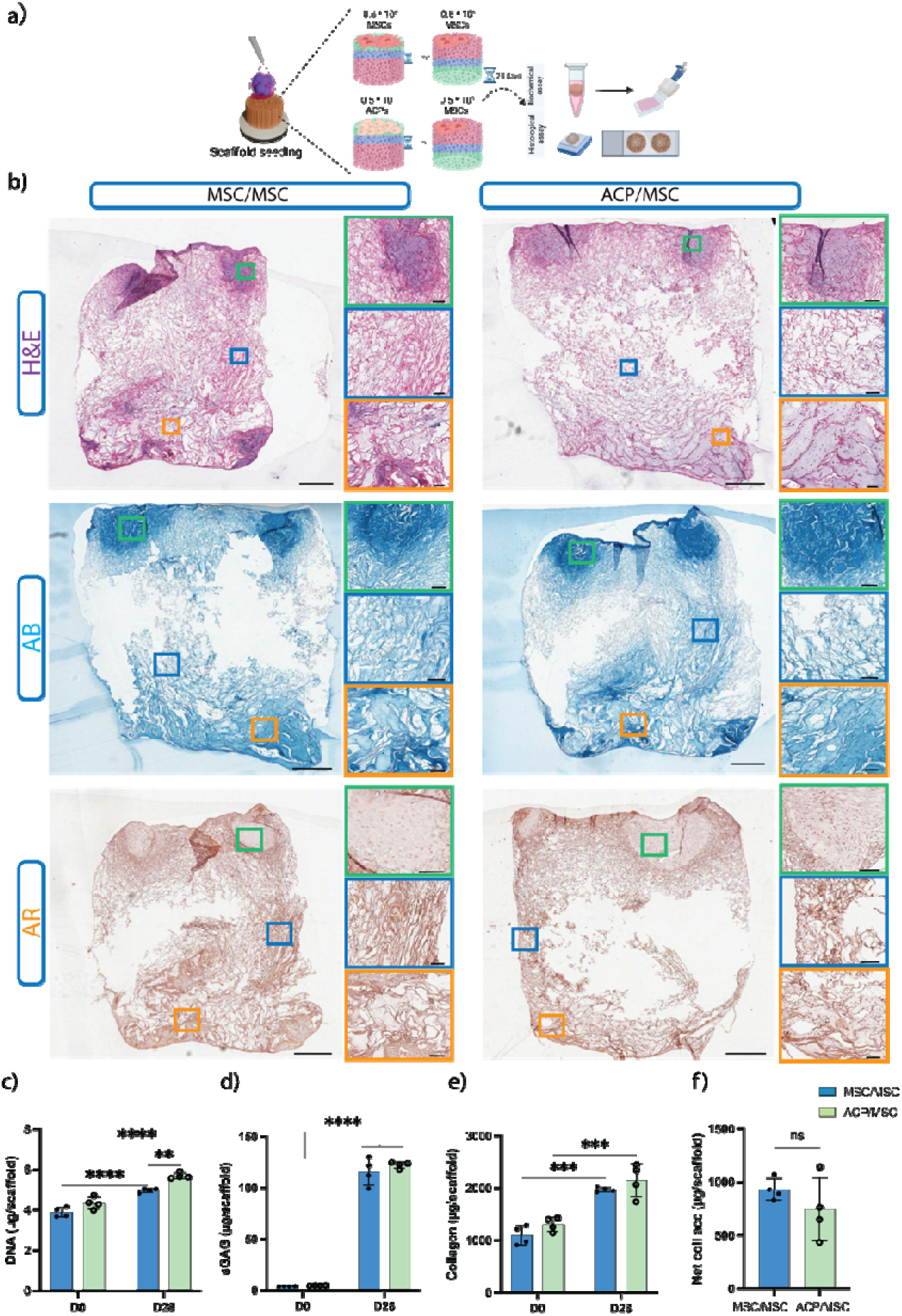
Multilayer tissue-specific ECM scaffolds support matrix deposition in vitro. (a) Schematic of scaffold seeding and analysis. In the MSC/MSC group, MSCs were seeded on both faces; in the ACP/MSC group, ACPs were seeded on the AC-ECM face and MSCs on the BN-ECM face. (b) Representative H&E, Alcian Blue and Alizarin Red staining after 28 days. (c–e) DNA, sGAG and total collagen content at days 0 and 28. (f) Net collagen accumulation, calculated as day-28 collagen minus day-0 collagen. Scale bars = 500 µm and 200 µm. Data are mean ± SD; n = 4 independent scaffolds. * p < 0.05, ** p < 0.01, *** p < 0.001, **** p < 0.0001.

A separate study was undertaken to assess the capacity of the multilayered ECM scaffolds to support region-specific tissue formation *in vitro* when seeded with only MSCs. Following 28 days of culture, immunohistochemical quantification showed regional differences in collagen deposition, with greater collagen type II staining in the AC region and greater collagen types I and X staining in the BN region (Supplementary Figure 1a, b).

### 3.3. Cell-seeded multilayer scaffolds promote spatially defined vascularisation and tissue formation in vivo

We next sought to determine the capacity of the multilayered scaffolds to support spatially defined vascularization and tissue formation *in vivo* following subcutaneous implantation into nude mice. Following 8 weeks *in vivo*, the multilayer scaffolds were analysed across three predefined regions corresponding to the chondral (2% AC), interface (5% AC) and osseous (5% BN) phases of the construct (Figure 3). MicroCT revealed limited mineral deposition within the middle and bottom regions of some MSC-only and co-culture scaffolds, whereas no mineral formation was observed in the unseeded scaffolds (Figure 3b). However, no significant differences in mineral deposition were detected between groups (Figure 3c). Histological analysis demonstrated tissue infiltration throughout the constructs, with significantly greater pore infiltration in both cell-seeded groups than in unseeded controls across all three regions (Figure 3d-f).

**Figure 3.**
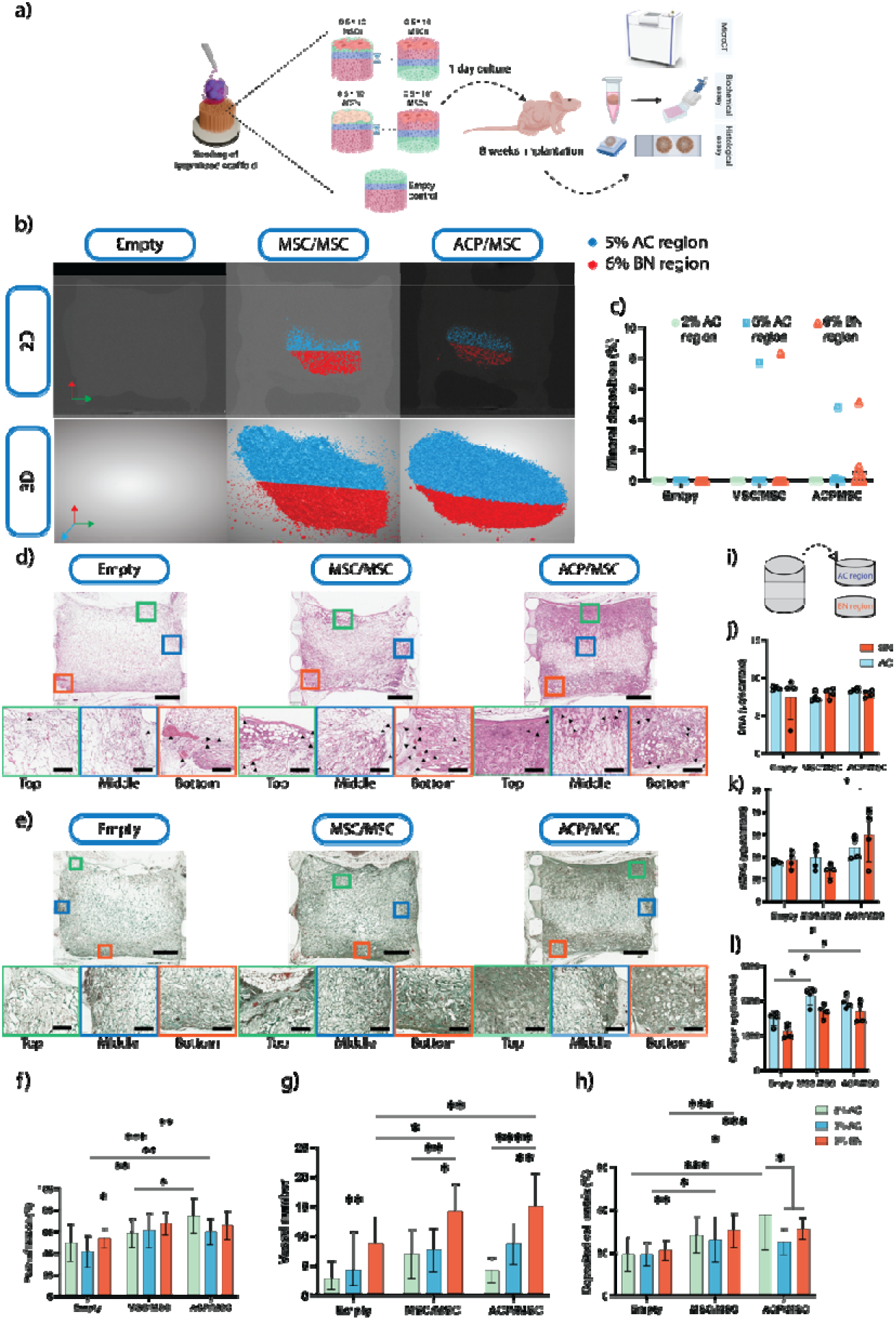
Cell-seeded multilayer scaffolds promote tissue infiltration, vascularisation and matrix deposition following subcutaneous implantation. (a) Study schematic. (b) Representative 2D and 3D µCT reconstructions of unseeded, MSC/MSC and ACP/MSC scaffolds after 8 weeks. Blue and red overlays show mineralised voxels assigned to the predefined 5% AC-ECM and 6% BN-ECM regions, respectively; these fixed regions reflect fabricated geometry and do not indicate sharply defined biological interfaces. (c) Mineral deposition in the three predefined regions. (d) H&E and (e) Masson’s trichrome staining. (f) Pore infiltration, (g) vessel number and (h) collagen-positive area. (i) Schematic of division into AC-ECM and BN-ECM regions for biochemical analysis. (j–l) DNA, sGAG and collagen content. Scale bars = 500 µm and 200 µm. Data are mean ± SD. For µCT, n = 8 scaffolds per group. After µCT, four scaffolds per group were allocated to histology/immunohistochemistry and four to biochemical analysis. Four sections were analysed per histological scaffold and averaged to provide one biological value per scaffold (n = 4). * p < 0.05, ** p < 0.01, *** p < 0.001.

Vessel desity was greatest within the bottom 6% BN region, reaching approximately 15 vessels per analysed ROI in both cell-seeded groups, compared with approximately 10 or fewer in the middle region and approximately 4 in the top region. Vessel number within the BN region was significantly greater in the MSC/MSC and ACP/MSC groups than in the corresponding region of unseeded scaffolds (Figure 3g). Masson’s trichrome staining demonstrated greater collagenous matrix deposition in cell-seeded constructs than unseeded controls (Figure 3e). Collagen-positive matrix occupied approximately 35–40% of the analysed area in both cell-seeded groups, compared with a maximum of ∼20% in unseeded scaffolds. The greatest collagenous matrix deposition was observed within the top region of the co-culture ACP/MSC group (Figure 3h).

For the biochemical analysis, the scaffolds were sectioned into a top chondral (AC) region and a bottom osseous (BN) region (Figure 3i). No significant differences in DNA content were observed between groups. Similarly, sGAG content was largely comparable, except for a significant increase in the ACP/MSC group relative to the BN region of the MSC/MSC group (Figure 3j, k). Collagen content was significantly greater in the AC and BN regions of both cell-seeded groups than in the corresponding regions of the unseeded controls (Figure 3l).

Qualitative immunohistochemical analysis further revealed stronger collagen type II staining within both the AC-ECM and BN-ECM regions of the cell-seeded scaffolds compared with the corresponding regions of the unseeded controls. The most abundant type II collagen deposition was observed in the AC region of the ACP/MSC group. Collagen types I and X were also detected within the cell-seeded constructs, although their staining appeared comparatively faint (Figure 4).

**Figure 4.**
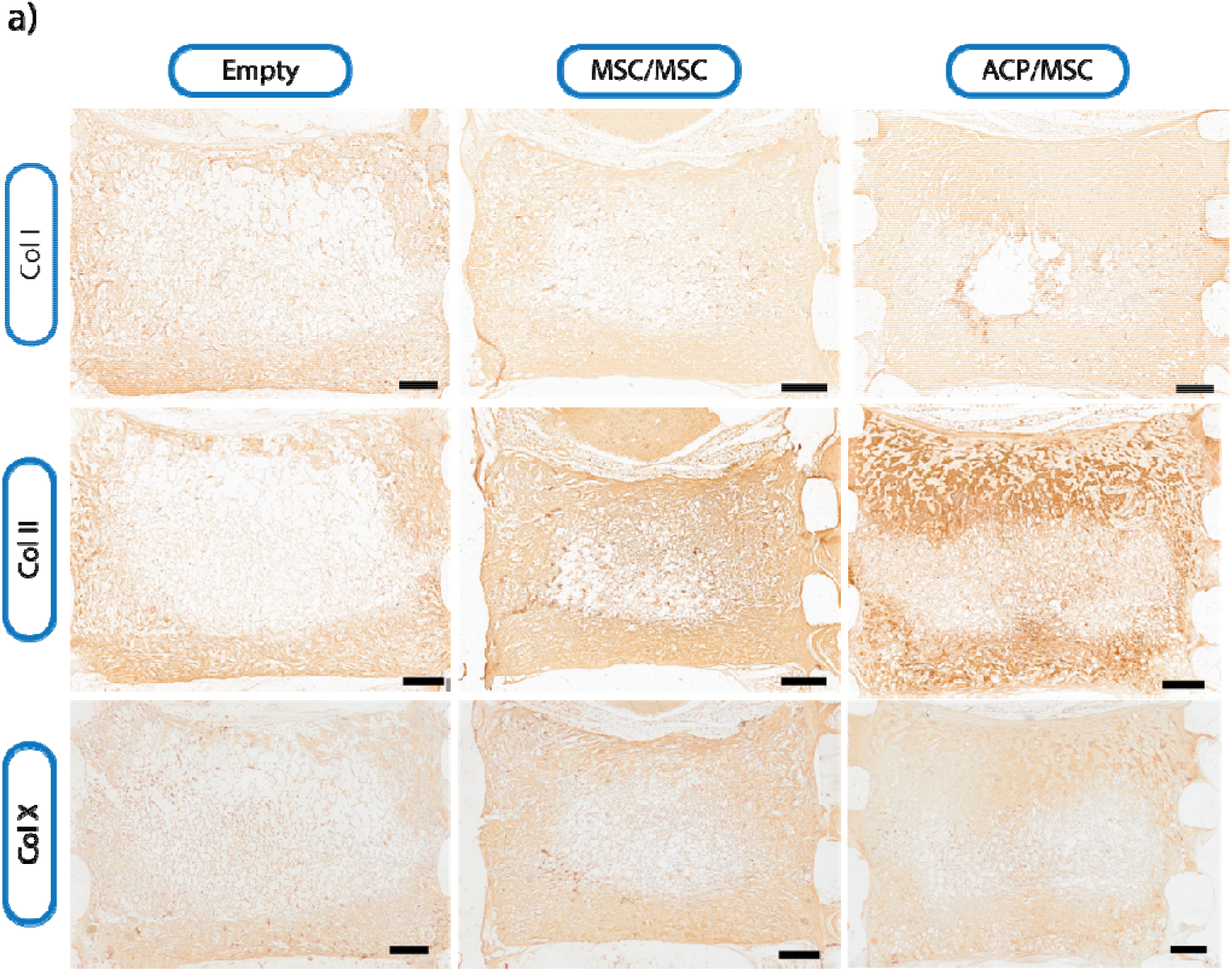
Collagen immunoreactivity within multilayer scaffolds following subcutaneous implantation. Representative immunohistochemical staining for collagen types I, II and X in unseeded, MSC/MSC and ACP/MSC scaffolds after 8 weeks. Brown staining indicates immunoreactivity and nuclei were counterstained with haematoxylin. Scale bar = 500 µm. Images are representative of n = 4 scaffolds per group.

### 3.4. Evaluation of osteochondral defect repair in a clinically relevant large animal model

The main goal of this study was to assess the capacity of acellular, multilayered ECM scaffolds to support osteochondral defect regeneration in a caprine large animal model. The surgical workflow is shown in Figure 5b. At six months, scaffold-treated defects had significantly higher ICRS macroscopic repair scores than paired empty defects (Figure 5c, d). Representative defects spanning the range of observed outcomes, including the best-, intermediate-, and worst-scoring defects from each group, are shown in Figure 5c. Note for the below analysis, the ‘best’, ‘intermediate’ and ‘worst’ healers were defined based on the combined results of post-explantation macroscopic scoring and an initial assessment of superficial articular cartilage repair using PTA-enhanced µCT.

**Figure 5.**
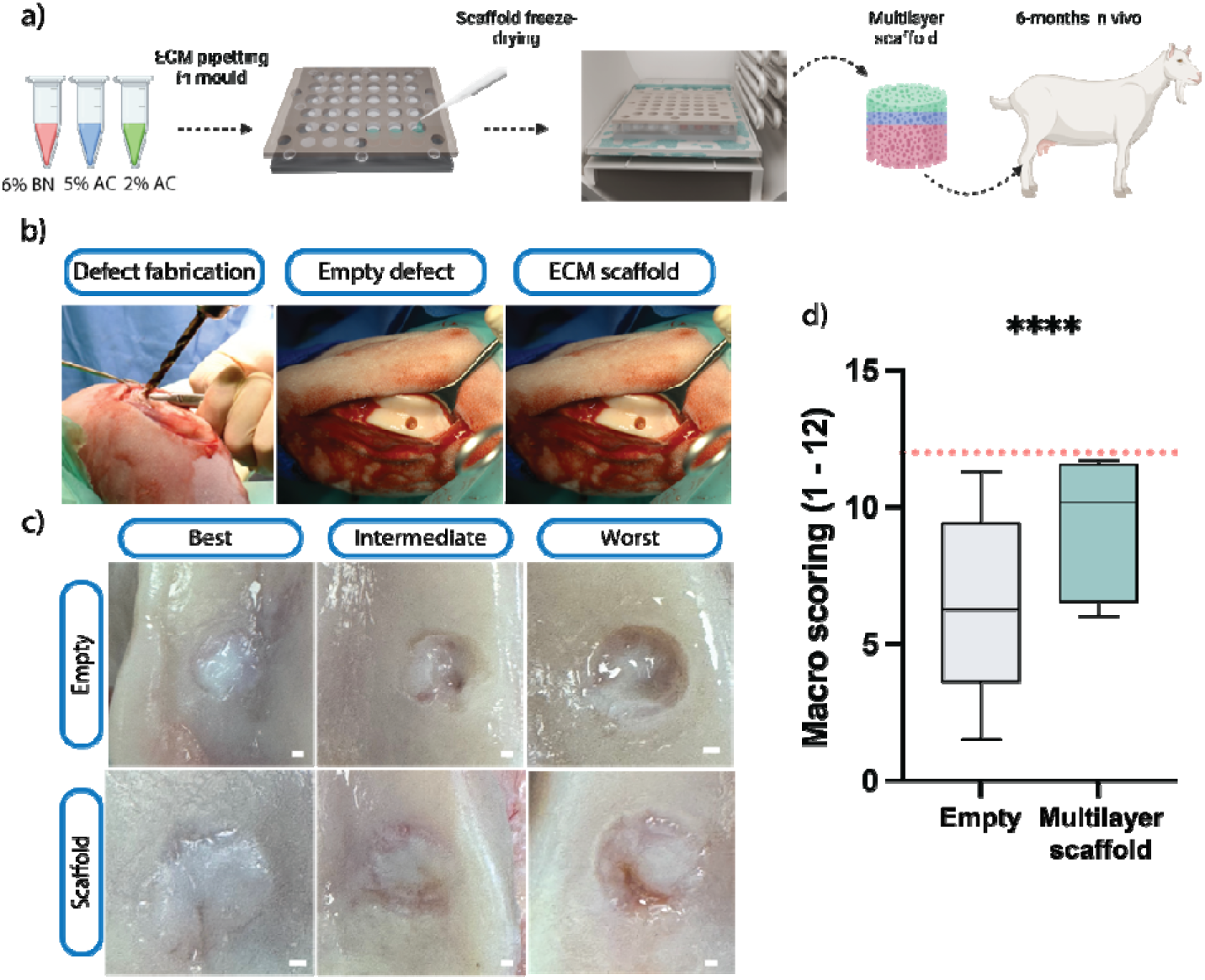
Multilayer ECM scaffold implantation improves macroscopic osteochondral defect repair after 6 months. (a) Caprine study design. (b) Surgical workflow showing creation of the 6-mm-diameter, 6-mm-deep defect and press-fit implantation of the 7-mm-diameter acellular scaffold. (c) Macroscopic appearance after 6 months. Representative best, intermediate and worst outcomes were selected according to the ICRS macroscopic score; the intermediate specimen was closest to the group median. (d) ICRS macroscopic repair scores. Scale bar = 1 mm. Box plots show median, interquartile range and full range; n = 10 paired defects per group. **** p < 0.0001. **[AUTHOR QUERY—FIGURE 5: Confirm the ranking criterion, intermediate-specimen rule and box-plot whisker definition used in the embedded graph.]**

Contrast-enhanced microCT analysis was used to determine both the extend of new bone formation in the osseous region of the defect, as well as amount of collagen in the chondral region of the defect using PTA which binds selectively to collagen (Figure 6a, b). This analysis confirmed improved cartilage regeneration in the scaffold treated defects, with a significantly higher collagen deposition in the chondral region of the scaffold treated group (approximately 60%) compared to the empty control (approximately 40%, Figure 6c). No significant difference was found in new tissue deposition in the transition (subchondral) and deep (bone) regions between groups (Figure 6d).

**Figure 6.**
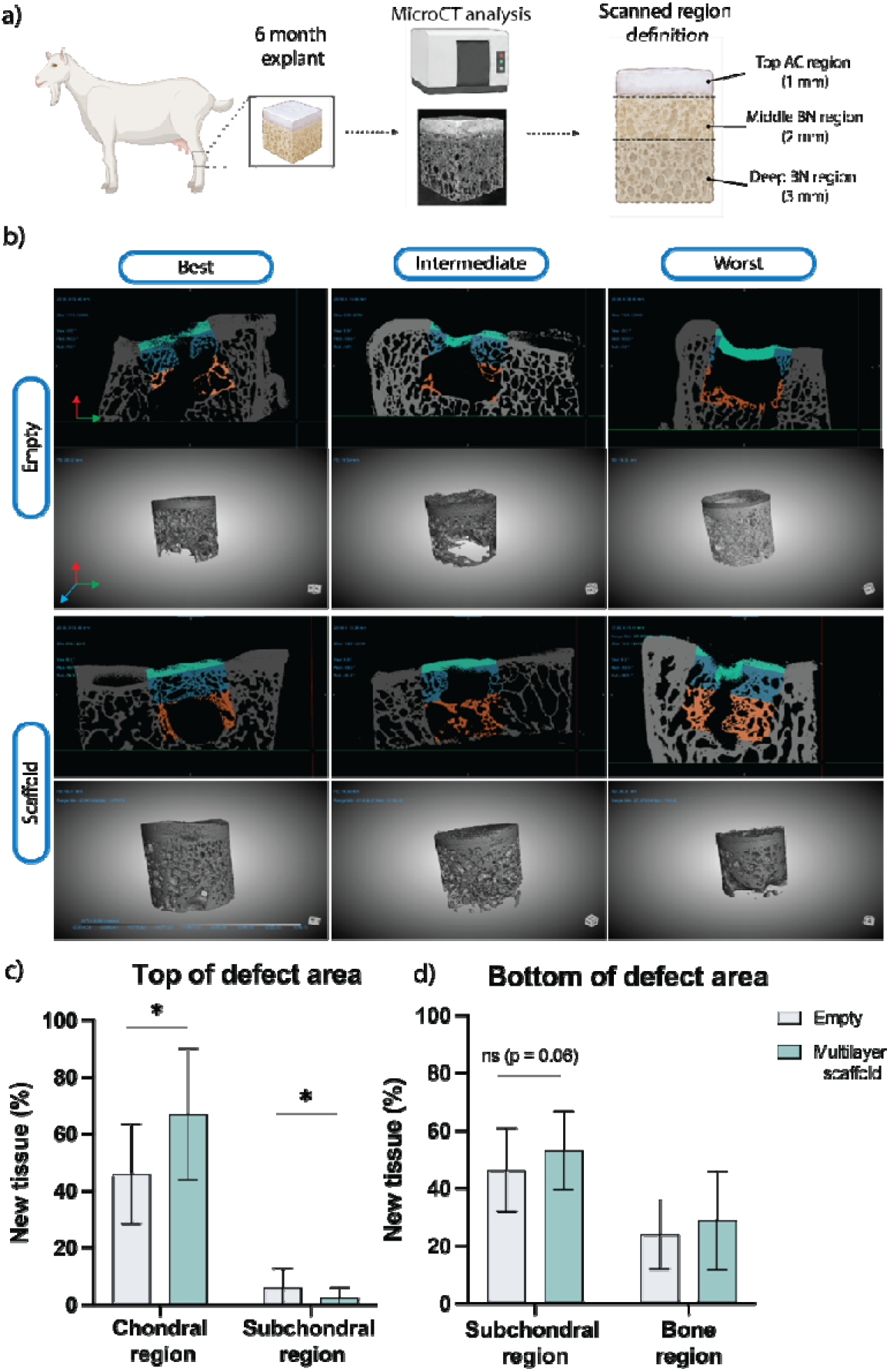
Contrast-enhanced µCT assessment of collagen-rich and mineralised repair tissue within caprine osteochondral defects. (a) Upper chondral (1 mm), middle subchondral (2 mm) and deep osseous (3 mm) regions of interest. (b) Representative reconstructions after six months. (c) PTA-attenuating collagen-rich repair-tissue fill within the upper chondral and middle subchondral regions. (d) Mineralised tissue fill within the middle subchondral and deep osseous regions. Coloured overlays are threshold-based segmentations and should not be interpreted as histologically confirmed cartilage or sharp biological interfaces. Data are mean ± SD; n = 10 paired defects per group. * p < 0.05.

Histological evaluation showed significantly higher ICRS II repair scores in the scaffold-treated group than in the empty control group (Figure 7c). Representative sections corresponding to the best-, intermediate-, and worst-scoring defects are shown for the empty and scaffold-treated groups in Figure 7a and Figure 7b, respectively. Polarised light microscopy identified differences in collagen fibre organisation within the superficial region of the repaired cartilage. The mean collagen fibre orientation in the scaffold-treated defects was significantly closer to 0° relative to the articular surface, while angular dispersion was significantly lower than in the empty defects (Figure 7d, e). Neither fibre orientation nor dispersion differed significantly between the scaffold-treated defects and native cartilage. In the deep cartilage region, fibre orientation and dispersion did not differ significantly among the native, scaffold-treated, and empty groups (Figure 7f, g).

**Figure 7.**
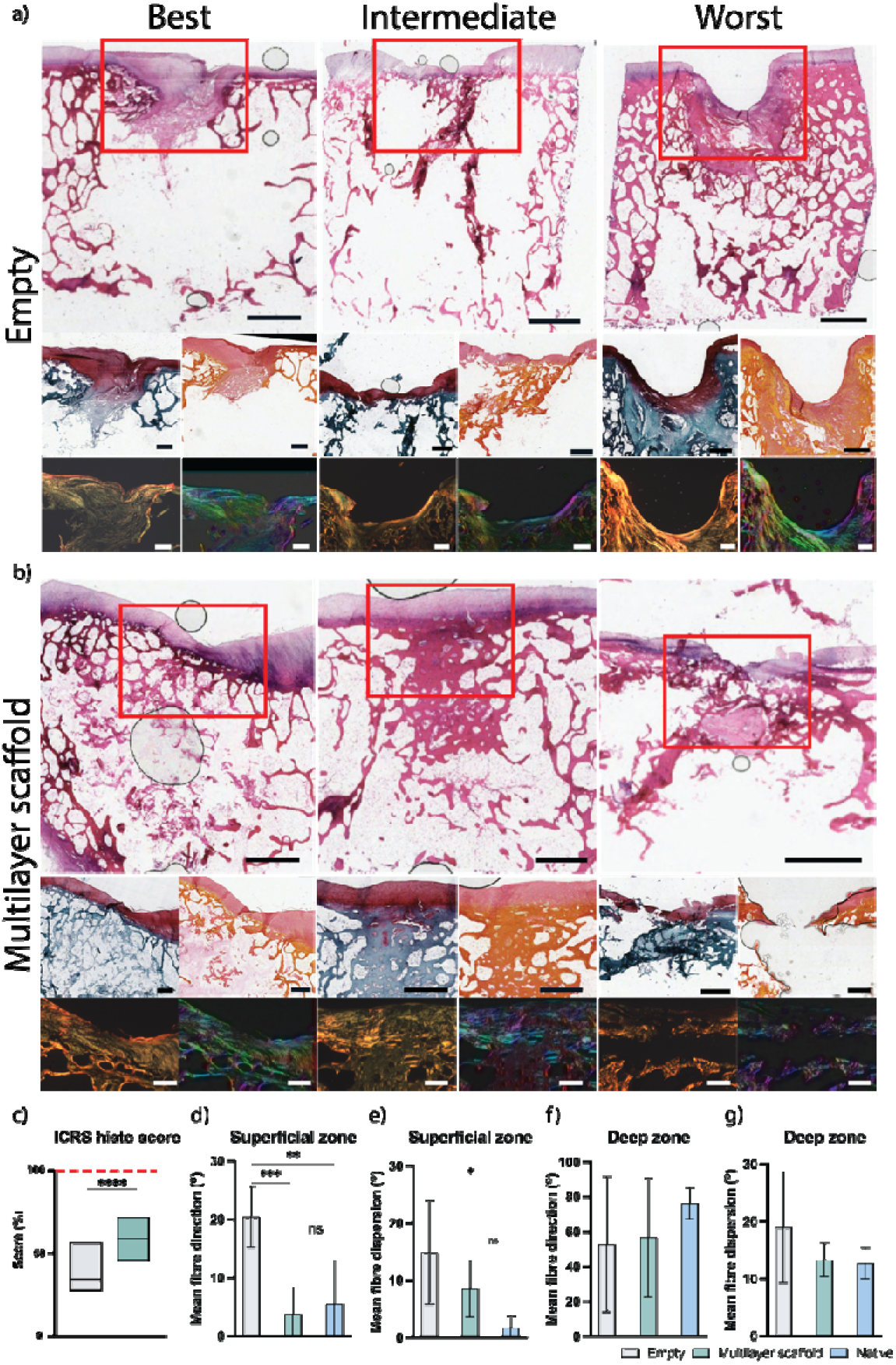
Multilayer ECM scaffolds improve histological repair and superficial collagen fibre organisation. (a,b) Representative histological and polarised-light images of empty and scaffold-treated defects after 6 months. Best, intermediate and worst outcomes were selected according to [CONFIRM: ICRS II histological score or the macroscopic ranking used in Figure 5]. (c) ICRS II repair scores. (d,e) Mean fibre orientation relative to the articular surface and angular dispersion in the superficial cartilage region. (f,g) Mean orientation and dispersion in the deep cartilage region. Native cartilage was included as reference. An orientation of 0° denotes fibres parallel to the articular surface; lower dispersion denotes more coherent alignment. Scale bars = 1 mm and 500 µm. In (c), box plots show median, interquartile range and full range. Data in (d–g) are mean ± SD. n = 10 defects per treatment group. * p < 0.05, ** p < 0.01, *** p < 0.001, **** p < 0.0001.

Immunohistochemical analysis revealed more intense collagen type II staining than collagen types I and X within the repair matrix of the scaffold-treated defects (Figure 8b). In contrast, the empty defects showed predominantly collagen types I and X staining, with positive collagen type II staining observed only in the highest-scoring empty defect (Figure 8a). RT-qPCR analysis of biopsies of the repair tissue confirmed this finding, with significantly higher expression of *ACAN* and *COL2A1* in the regenerated articular cartilage of scaffold-treated defects compared with empty controls (Figure 8c, d). Conversely, *MMP13* and *COL10A1* expression was significantly lower in the scaffold-treated defects (Figure 8e, f).

**Figure 8.**
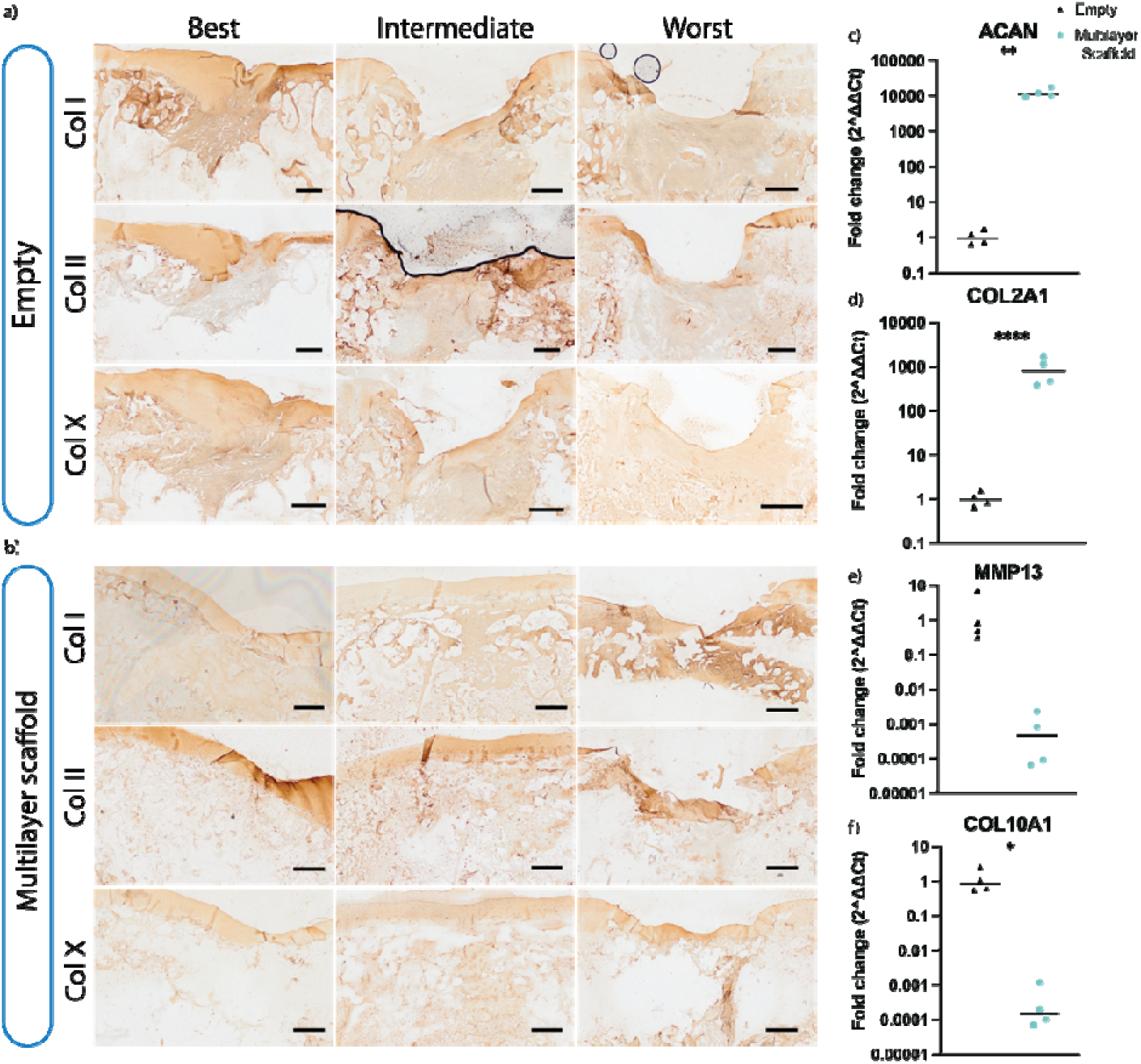
Multilayer ECM scaffolds promote a more chondrogenic repair-tissue phenotype. (a,b) Representative immunohistochemical staining for collagen types I, II and X in empty and scaffold-treated defects. Best, intermediate and worst outcomes were selected according to [CONFIRM: ICRS II histological score or the macroscopic ranking used in Figure 5]. Brown staining indicates immunoreactivity. (c–f) RT-qPCR analysis of *ACAN*, *COL2A1*, *MMP13* and *COL10A1* in repair tissue from the articular cartilage region. Expression was normalised to 18S rRNA and expressed relative to the paired empty control using the 2^−ΔΔCt method. Each symbol represents one animal and the horizontal line denotes the mean; n = 4 paired animals. Scale bar = 1 mm. * p < 0.05, ** p < 0.01, **** p < 0.0001.

## 4. Discussion

The overall goal of this study was to develop an “off-the-shelf”, multilayered ECM-derived scaffold that recapitulates key aspects of the spatial complexity of the osteochondral unit and promotes region-specific tissue formation. To achieve this, cartilage- and bone-derived ECM biomaterials were integrated through a sequential freeze-drying strategy to generate three continuous yet structurally distinct regions with highly elastic mechanical properties. We demonstrated that these region-specific microenvironments supported cell survival and spatially directed differentiation *in vitro* and vascularization and tissue formation *in vivo* following subcutaneous implantation. Finally, implantation in a clinically relevant caprine osteochondral defect model improved repair compared with untreated defects, promoting the development of more structurally organised hyaline cartilage.

The continuous structure of the multilayer scaffold indicates that ECM concentration and tissue source could be varied without creating mechanically weak boundaries between phases. A similarly integrated architecture was reported by Levingstone et al. [51], who developed a multilayer collagen-based scaffold in which gradual changes in composition reduced the risk of delamination between the cartilage and bone regions. In the present study, combining the three ECM phases also produced a construct with excellent shape recovery following repeated compression, despite the different mechanical properties of the individual layers of the scaffold. Of note is that the scaffold is considerable softer than native osteochondral tissue, and future studies should determine whether it can maintain its regenerative properties when used to treat larger osteochondral defects, which may present a more mechanically challenging environment.

When seeded with only MSCs, the chondral region of the construct preferentially supported the deposition of sGAGs and type II collagen *in vitro*, suggesting the retention of tissue-specific signals throughout the depth of the multilayered scaffold following ECM processing. This agrees with previous studies [47,52], where it was demonstrated that scaffolds containing articular cartilage and growth-plate ECM could spatially direct stable chondrogenesis and endochondral ossification, respectively. Unlike the particulated ECM used in those studies, the ECM employed here underwent solubilisation before scaffold fabrication. The maintenance of regional phenotypes with scaffold depth therefore suggests that sufficient compositional or structural cues remained after processing to influence MSC differentiation.

Following subcutaneous implantation, the cell-seeded multilayer scaffolds promoted spatially defined vascularization and tissue formation. Vascular invasion in vivo was highest within the BN-ECM region of the multilayered scaffold, suggesting that either the AC ECM is suppressing vascularization, and/or that the BM-ECM contains factors supportive of angiogenesis. It is well established that the cartilage matrix contains factor known to suppress vascularization, including thrombospondins, chondromodulin-I and endostatins [53], and we have previously shown that AC ECM scaffolds supress angiongenesis *in vivo* compared to other ECM types [26]. We have also previously shown that BN-ECM, and hypertrophic cartilage, is supportive of angiogenesis *in vivo* [22,54,55]. It should be noted, however, that the increased levels of vascularisation observed here in the BN region of the multilayered scaffold was not accompanied by substantial mineral formation. This suggests that vascular infiltration alone was insufficient to initiate robust ossification in a subcutaneous environment. In the same *in vivo* environment, the cell-seeded scaffolds supported higher levels of collagen type II deposition than collagen types I and X, particularly in the chondral region of the construct, consistent with the development of a hyaline cartilage-like tissue.

In the goat model, the clearest effect of the multilayered scaffold was observed within the cartilage region of the osteochondral defect. This is important because untreated critical-sized osteochondral defects in goats generally demonstrate poor and inconsistent restoration of the articular surface [56]. Furthermore, previous studies showed progressive osteochondral repair using a cell-free multilayer collagen scaffold, although the cartilage remained relatively immature at six months [29,57]. In this study, the improved macroscopic and histological repair observed at six months therefore indicates that our multilayered ECM supported hyaline cartilage formation at a comparatively early stage of repair. Further studies are required to assess if this can be maintained at later timepoints.

Polarised light microscopy provided additional evidence that the scaffold influenced the quality and not only the amount of repair tissue. Although previous studies show how introducing aligned pores into a lyophilised ECM scaffold can direct collagen organisation [29], the present scaffold did not contain a deliberately aligned superficial pore network, yet the collagen fibres within the superficial repair tissue adopted a more native-like orientation than those in empty defects. This may indicate that the AC-ECM environment supported matrix maturation while joint loading subsequently directed collagen fibre organisation. The predominance of collagen type II, together with increased chondrogenic and reduced hypertrophic and catabolic gene expression, further supports the formation of a more stable hyaline cartilage-like matrix, which contrasts with the type I-rich fibrocartilage commonly reported following unsuccessful cartilage repair [58].

The main limitation of the large-animal study is the comparison of the multilayer scaffold with only an empty defect. Future studies should include single-phase ECM scaffold controls, together with a clinically relevant comparator. Evaluation at only one endpoint also prevented assessment of scaffold degradation, early-stage immune response and the progression of tissue maturation. Finally, explant mechanical testing and more detailed analysis of subchondral bone architecture will be required to determine whether the improved structural repair translates into functional and durable osteochondral regeneration.

## 5. Conclusion

Multilayered tissue-specific ECM support region-dependent cellular differentiation *in vitro* and spatially defined vascularization and tissue development *in vivo*. In a caprine osteochondral defect model, scaffold treatment improved macroscopic and histological repair, increased tissue formation within the cartilage region, restored a more native-like superficial collagen organisation and promoted the formation a collagen type II-rich articular cartilage with reduced expression of hypertrophic and catabolic markers compared to empty controls. Although further work is required to establish functional restoration, long-term durability and improved subchondral bone regeneration, these findings support the continued use of multilayered ECM scaffold for osteochondral repair.

## CRediT authorship contribution statement

Giovanni Gonnella: Conceptualization, Methodology, Investigation, Formal analysis, Writing – original draft, Writing – review & editing.

Olivia Strong: Methodology, Formal analysis.

Vasile M. Sularea: Methodology.

Gabriela S. Kronemberger: Methodology.

Aliaa S. Karam: Methodology.

Daniel J. Kelly: Conceptualization, Supervision, Writing – review & editing, Funding acquisition.

## Declaration of generative AI and AI-assisted technologies in the manuscript preparation process

During the preparation of this work, the authors used ChatGPT (OpenAI) to improve language and readability. After using this tool, the authors reviewed and edited the content as needed and take full responsibility for the content of the published article.

## Declaration of competing interests

Giovanni Gonnella and Daniel J. Kelly are named inventors on a patent application related to the scaffold technology described in this work. The authors declare no other competing financial or non-financial interests.

## Data Availability Statement

The processed data supporting the findings of this study are provided in the article and its Supporting Information. The underlying raw data are available from the corresponding author upon reasonable request.

## Acknowledgments

Schematic elements were created by G.G. using BioRender.com, Autodesk Fusion 360, Blender and Adobe Illustrator.

## Funding

This publication has emanated from research supported in part by a grant from Research Ireland under Grant No. 12/RC/2278_P2. This project received funding from the European Research Council under the European Union’s Horizon Europe research and innovation programme (4D-BOUNDARIES; Grant Agreement No. 101019344).

## Notes

### Summary of Updates

The new version modifies different parts of the manuscript, as well as removing one of the supplementary figures and correcting different grammatical errors.

