## Supplementary information for "Multilayered extracellular matrix–derived scaffolds direct progenitor cell differentiation *in vitro* and osteochondral-tissue formation *in vivo*"

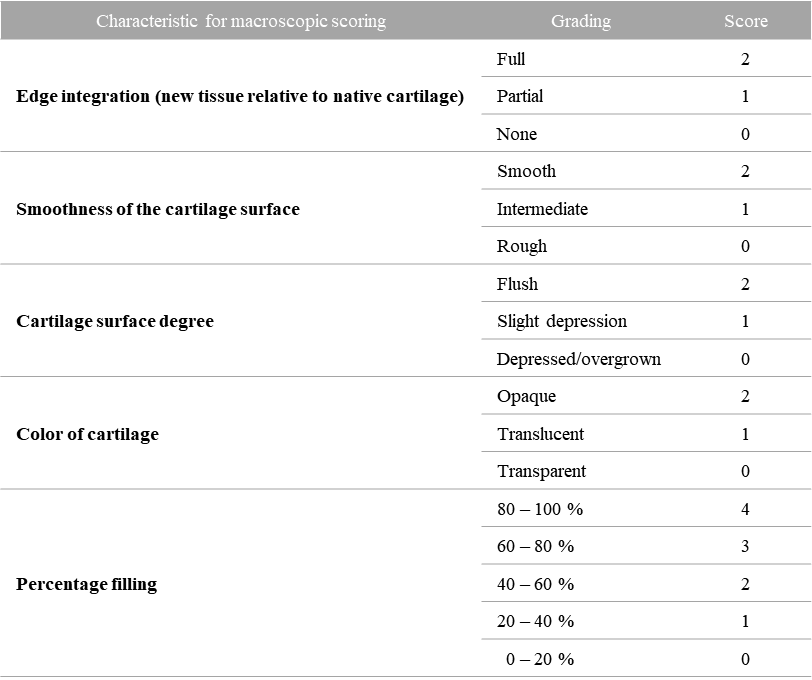

**Supplementary Table 1. Macroscopic scoring system for cartilage repair. Maximum score possible is 12.**

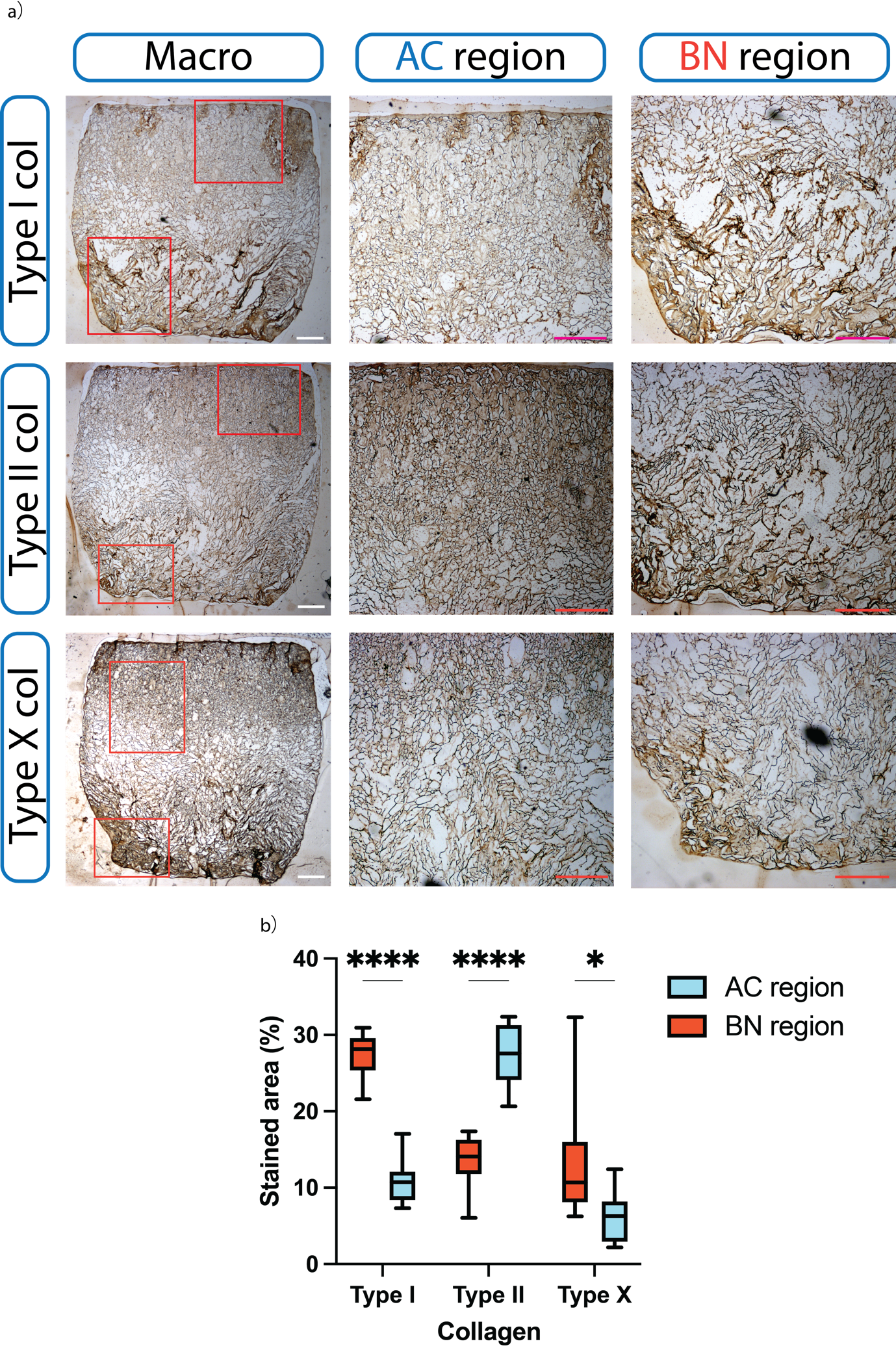

**Supplementary Figure 1. Regional collagen phenotype within enlarged multilayer ECM scaffolds cultured with goat MSCs.** (a) Representative immunohistochemical staining for collagen types I, II and X within the AC and BN regions following 28 days of culture. Brown staining indicates immunopositivity. (b) Quantification of the percentage positively stained area for each collagen type within the AC and BN regions. Scale bar = 1 mm, 500 µm. Data are presented as mean ± SD. n = 4, * p < 0.05, **** p < 0.0001.
